# A low-cost platform suitable for sequencing-based recovery of natural variation in understudied plants

**DOI:** 10.1101/2020.06.24.169276

**Authors:** Rachel Howard-Till, Claudia E. Osorio, Bradley J. Till

## Abstract

Genetic characterization of wild and cultivated plants provides valuable knowledge for conservation and agriculture. DNA sequencing technologies are improving and costs are dropping. Yet, analysis of many species is hindered because they grow in regions that lack infrastructure for advanced molecular biology. We developed and adapted low-cost methods that address these issues. Tissue is collected and stored in silica-gel, avoiding the need for liquid nitrogen and freezers. We have optimized low-cost home-made DNA extraction to increase yields, reduce costs, and produce DNA suitable for next generation sequencing. We also describe how to build a gel documentation system for DNA quantification. As a proof of principle, we use these methods to evaluate wild *Berberis darwinii*, native to Southern Chile.

**Method summary:** We describe a suite of low-cost do-it-yourself methods for field collection of plant tissues, extraction of genomic DNA suitable for next generation sequencing, and home-made agarose gel documentation suitable for DNA quantification. These methods enable the collection and preparation of samples for genomic analysis in regions with limited infrastructure.

---

Although plants have been domesticated for more than 10,000 years, just twelve crops make up 75% of the world’s plant-based foods [1]. In these, genetic erosion has greatly reduced sequence diversity, limiting efforts to adapt them to changing environments [2–4]. However, there are more than 10,000 edible species worldwide, where allelic diversity provides resilience to biotic and abiotic stresses and productivity in poor soils [5]. These plants are the foundation for sustainable agriculture accomplished through precision domestication [6]. Cataloging genetic diversity is an important step in realizing the genetic potential of new crops, and also to guide conservation of wild species. While tremendous advances in DNA sequencing have been reported, a bottleneck exists in that many promising species are often growing in regions lacking advanced laboratories. While genomic DNA can be shipped to a sequencing facility, researchers need robust, cost-effective on-site methods for field collection of tissue, extraction of genomic DNA, and subsequent measurement of quantity and quality of DNA.

We have developed a low-cost platform suitable for reduced representation sequencing of understudied plants, where each step of the procedure can be implemented in laboratories with limited infrastructure. We used *Berberis darwinii*, a native berry from Southern Chile, as a proof of principle. The first step in the process is collection of plant tissues. A common method to preserve genomic DNA uses immersion of fresh tissue in liquid nitrogen. The disadvantage of this approach is that it requires liquid nitrogen and −80°C freezers to store tissue. Lyophilization is a suitable alternative, but this requires specialized equipment [7,8]. An easy, low-cost alternative is collection and storage of tissue on silica gel at room temperature [9,10]. This approach was previously shown to be suitable for preparation of genomic DNA of suitable quality for PCR and agarose gel-based assays [11]. We modified this method for field-collection of leaf samples from hundreds of plants. Leaf tissues are collected in tubes, covered with silica-gel and stored for at least two days until tissues are desiccated (Figure 1, for detailed protocol see dx.doi.org/10.17504/protocols.io.bdg9i3z6). This method is highly portable, low-cost (silica-gel is reusable), and tissues can be stored in silica-gel for months to years.

**Figure 1:**
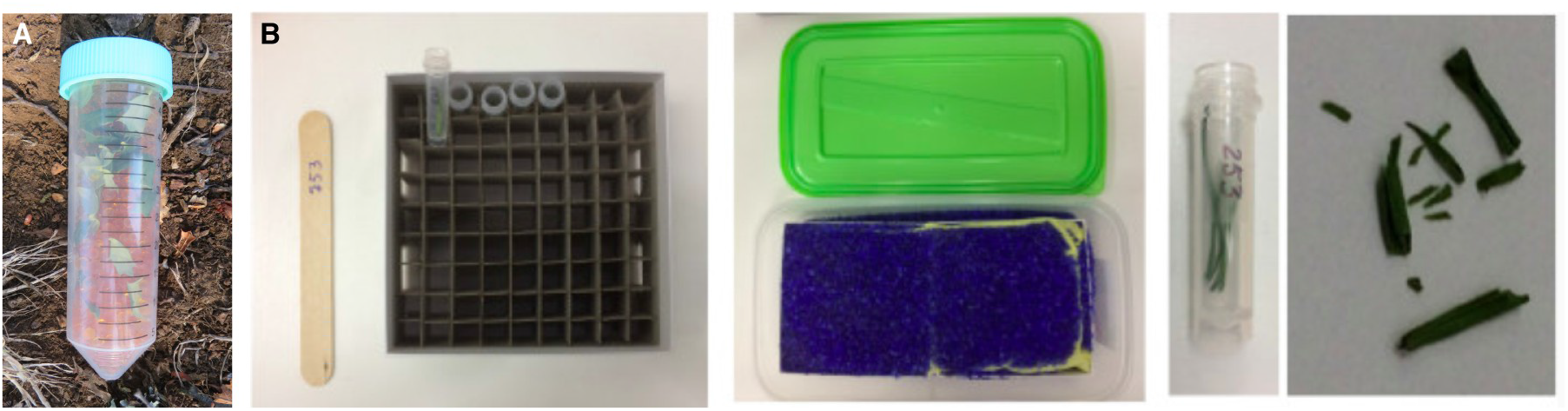
Plant tissues collected in silica gel for desiccation. A. Example of collection of *Berberis darwinii* leaves in a 50 ml tube. The size and shape of *B. darwinii* leaves makes placement into 2 ml tubes impractical. Multiple leaves (approximately 30) are collected from a single plant. B. Collection of flax leaves in 2 ml tubes. To facilitate field collection, wooden stakes and tubes are labeled with plant numbers prior to collection. Tissue is collected from field-grown material into two boxes (approx. 200 samples) before applying silica. Tube boxes are placed in a plastic container then covered with a porous material such as a paper towel and covered in silica gel and then sealed (second panel, silica gel with blue indicator is used in this example). Tissue shrinks when dry (third panel) and is brittle when fully desiccated (fourth panel).

The next step is extraction of genomic DNA. Methods using organic-phase separation can produce high quality DNA suitable for next generation sequencing [12,13]. However, this requires procurement, safe storage, use, and disposal of organic chemicals. Commercial DNA kits, while more costly, avoid the use of toxic organic-phase separation and instead use chaotropic salts and silica matrix binding. We have previously described a method using self-prepared buffers and silica matrix that provides high quality DNA at approximately one tenth the cost of a kit [14]. We modified this protocol to increase DNA yields because larger quantities of DNA are required for reduced representation sequencing. The full protocol is available at https://dx.doi.org/10.17504/. For comparison, one replicate sample was prepared using the DNeasy Plant Mini kit, (Qiagen, Germantown, MD, USA), with a slight modification of the lysis step because it was observed that lysis at 65° C as specified in the provided protocol caused the DNA to be degraded. This step was therefore performed at room temperature and no degradation was observed (Figure 2A).

**Figure 2.**
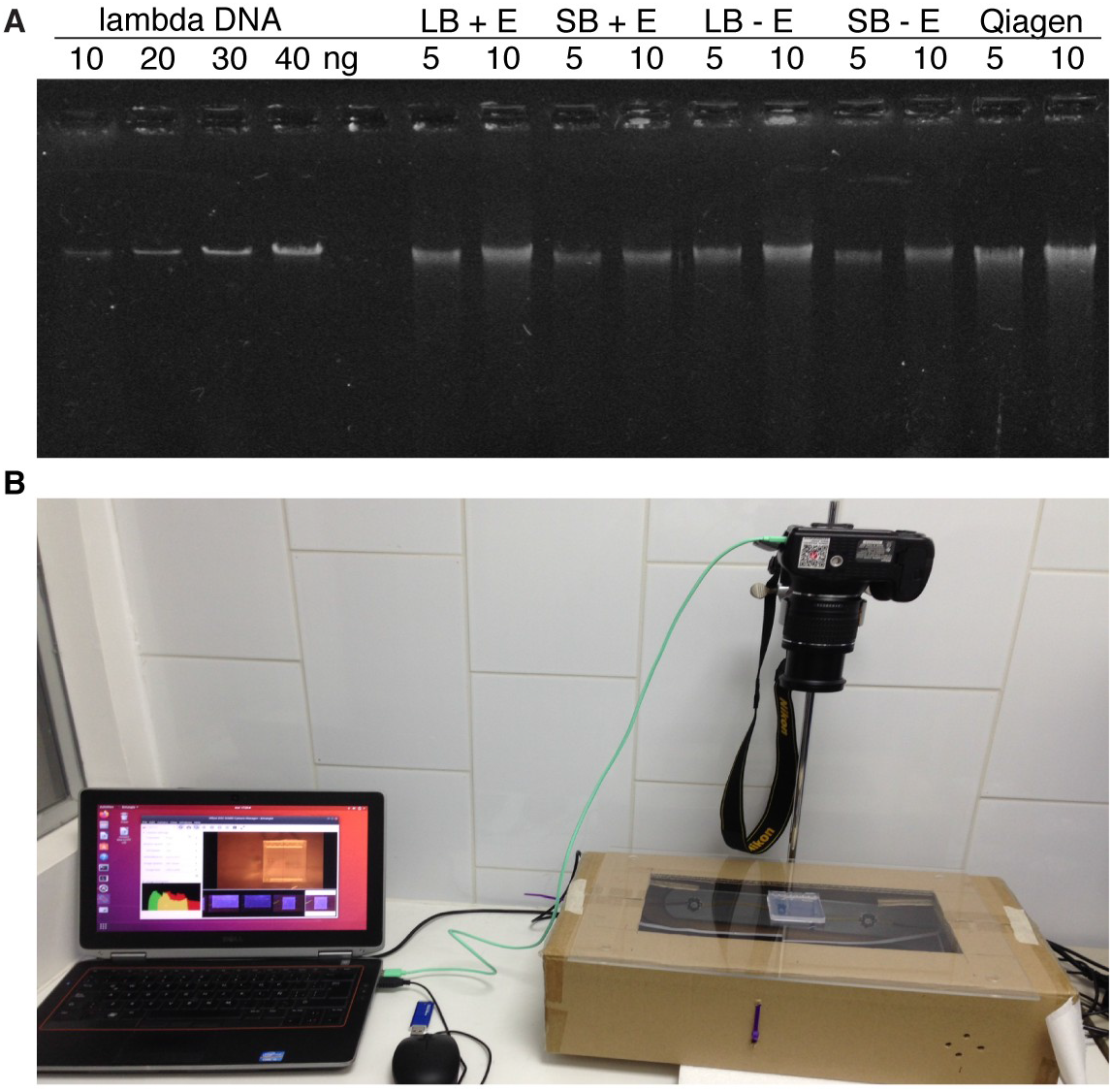
A, Low cost DNA extraction methods produce quantity and quality of genomic DNAs comparable to a commercial kit. 5 and 10 μl aliquots of eluted DNA samples prepared by low cost methods or a commercial kit were run on a 1% agarose gel and stained with SYBR safe DNA stain. Lambda DNA samples were included on the gel as mass standards for estimation of DNA concentration. LB, long binding time, SB, short binding time, +E, ethanol included in binding buffer, −E, no ethanol in binding buffer. B, Digital images of stained agarose gels are produced with a home-made gel documentation system consisting of a digital camera, a computer to control the camera and store images, and a light box created using blue LED lights.

After extraction, it is important to evaluate the quality and concentration of genomic DNAs by gel electrophoresis. However, equipment for image acquisition and analysis of agarose gels can cost tens of thousands of dollars. The development of DNA stains that fluoresce under blue lights allows the use of a gel documentation system consisting of a self-built blue-LED light box and a digital camera. Various tutorials for light box construction are available online, but many are highly technical (for example, https://openwetware.org/wiki/DIY_Blue_Gel_Transilluminator). We built a simple gel documentation system composed of low-cost electronic components and free software for less than 800 US dollars (for details and costs see: dx.doi.org/10.17504/protocols.io.bdtpi6mn). The system includes a blue LED light box and a Nikon D3400 digital camera, controlled using the free Entangle software and a recycled laptop running Linux Ubuntu (Figure 2B). This was used to evaluate the quality and concentration of *Berberis* genomic DNA samples by measuring the intensity of bands in gel images and comparing to lambda DNA standards of known concentration with the software ImageJ (Figure 2A and Table 1).

**Table 1:** Concentration and yield of genomic DNAs.

| Prep method | [eluted DNA] | Est. yield (µg) – gel | Yield (µg) – Qubit | ng DNA/mg dry tissue |
| --- | --- | --- | --- | --- |
| Long bind + EtOH | 11.83 | 5.9 | 4.93 | 41.0 |
| Short bind + EtOH | 7.67 | 3.8 | 3.02 | 27.5 |
| Long bind | 8.84 | 4.4 | 4.16 | 41.6 |
| Short bind | 6.65 | 3.3 | 2.62 | 23.8 |
| Qiagen kit | 14.01 | 3.9 | 6.97 | 69.7 |

Due to the large volume used for elution of genomic DNA, samples were concentrated by precipitating in 13.3% (final concentration) PEG 6000, 10 mM MgCl_2_ and resuspended in a final volume of 100 μl TE. To test the accuracy of the ImageJ quantification, DNA concentrations were also measured using a Qubit fluorometer and the dsDNA High Sensitivity assay (Thermo Fisher Scientific, Waltham, MA, USA). The maximum yield from the low-cost method was slightly less than that of the commercial kit, but overall the quality and quantity of DNA was acceptable for the purpose (Figure 2B and Table 1). DNA yield was improved by increasing binding time, but adding EtOH in the binding buffer did not significantly affect yield. However, using EtOH reduces the cost by reducing the volume of chaotropic buffer, the most expensive component of the assay. The adjusted cost is 32 cents (USD) per sample versus 42 cents without ethanol (calculated from [14]). Estimated yields from the Qubit assay were slightly lower than from the gel image, which may reflect a slight loss of DNA during the precipitation step, or may reflect slight differences in the methods. However, the measurements were generally proportional, with the exception of the Qiagen sample. The concentration of this sample may have been underestimated in the gel measurement due to the DNA being slightly less intact, causing more of the DNA to be outside the region of the gel image selected for quantification.

To prepare the DNA for reduced-representation genome sequencing, restriction enzyme(s) and a molecular weight range of DNA fragments are chosen to produce the desired number of unique base pairs. Because there is no reference genome for *Berberis darwinii*, the closely related *B. thunbergii* genome and the Perl module RestrictionDigest was used to determine the minimum amount of starting DNA needed to produce 200 ng for library preparation (see dx.doi.org/10.17504/protocols.io.bdtji6kn for detailed protocols for experimental design and post-sequencing analysis). 5 μg of genomic DNA was digested with 2 μl FastDigest BsuRI (an isoschizomer of HaeIII),(Thermo Fisher Scientific, Waltham, MA, USA) for 2 h at 37°C. Digested DNA was run on a 1% agarose gel in ½X TBE (Figure 3). The gel was stained with SYBR safe (Thermo Fisher Scientific, Waltham, MA, USA) and DNA fragments of between ~350 and 700 base pairs were excised from the gel and purified using the E.Z.N.A gel extraction kit (Omega bio-tek, Norcross GA, USA). DNA recovery was determined by Qubit assay (Thermo Fisher Scientific, Waltham, MA, USA). DNA was sent to Novogene (Novogene, Co., Ltd., Beijing, China) for library preparation and 2×150 PE sequencing on an Illumina Novaseq.

**Figure 3.**
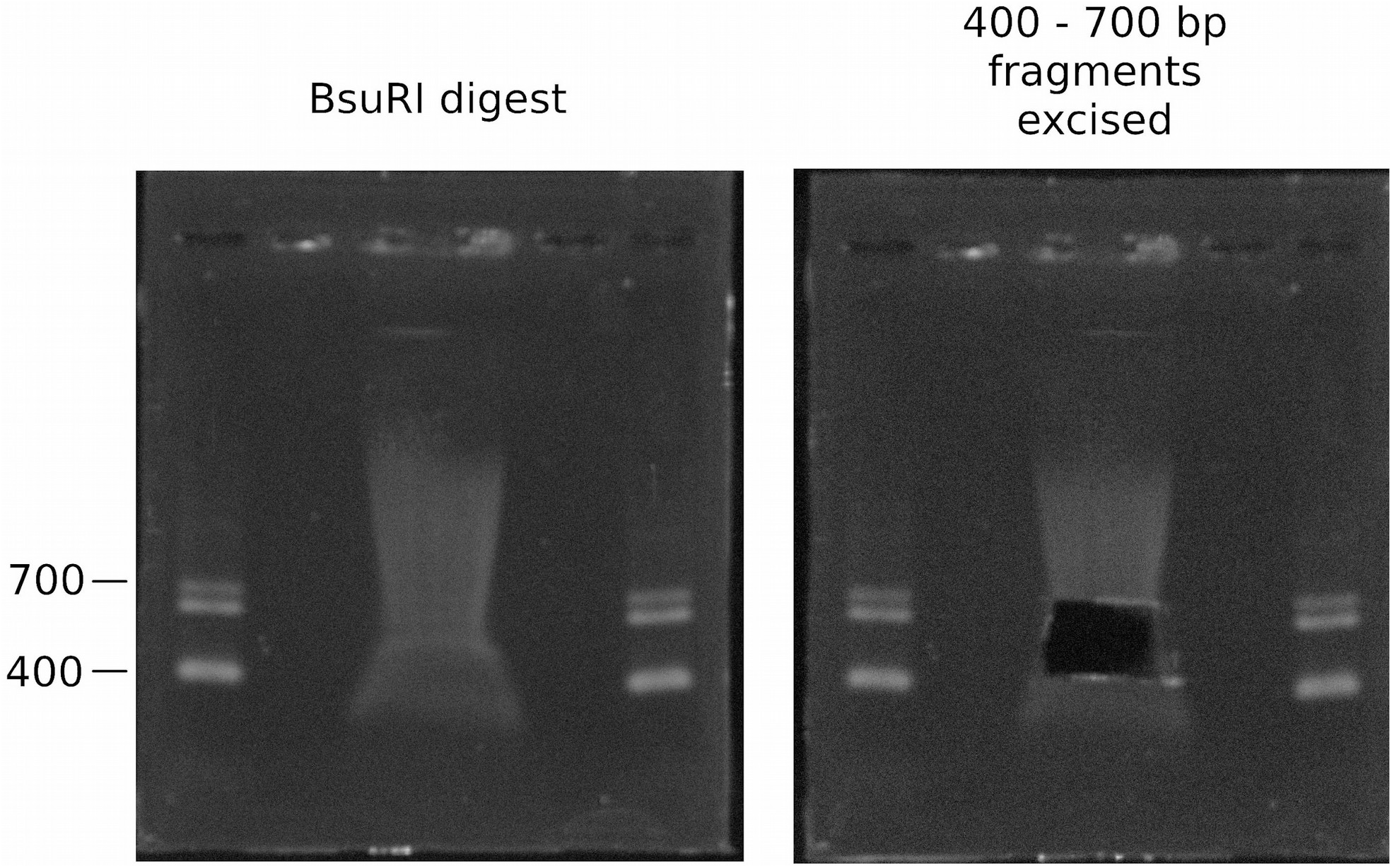
Isolation of digested DNA photographed on the home-made gel-doc.

Compressed fastq files were supplied by the sequence provider. FastQC analysis revealed high quality data produced from the genomic DNA using the low-cost methods (Figure 4). Fastq data was pre-sprocessed using the process radtags function in Stacks and interleaved using BBmap. Analysis proceeded using the Stacks *de novo* reference independent pipeline to produce BAM, VCF and fasta-formatted sequence files for the wild-type *Berberi*s sample. This resulted in a mean coverage of 8.7x, with 84 million unique base pairs producing more than 25,000 candidate heterozygous SNP polymorphisms (Figure 5, and dx.doi.org/10.17504/protocols.io.bdtji6kn; fastq data is available at https://www.ncbi.nlm.nih.gov/sra/?term=PRJNA613471).

**Figure 4:**
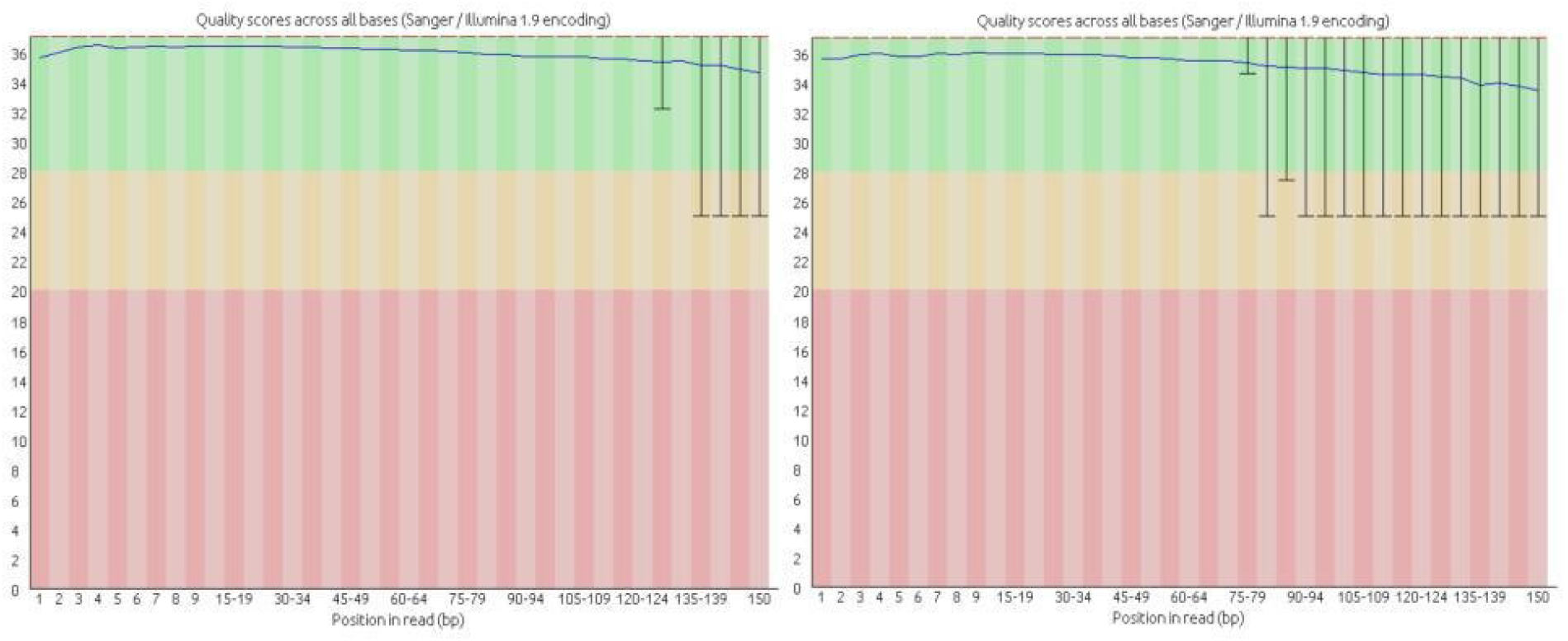
FastQC output of quality scores across all bases for read 1(left) and read 2 (right) of the 2×150PE run.

**Figure 5:**
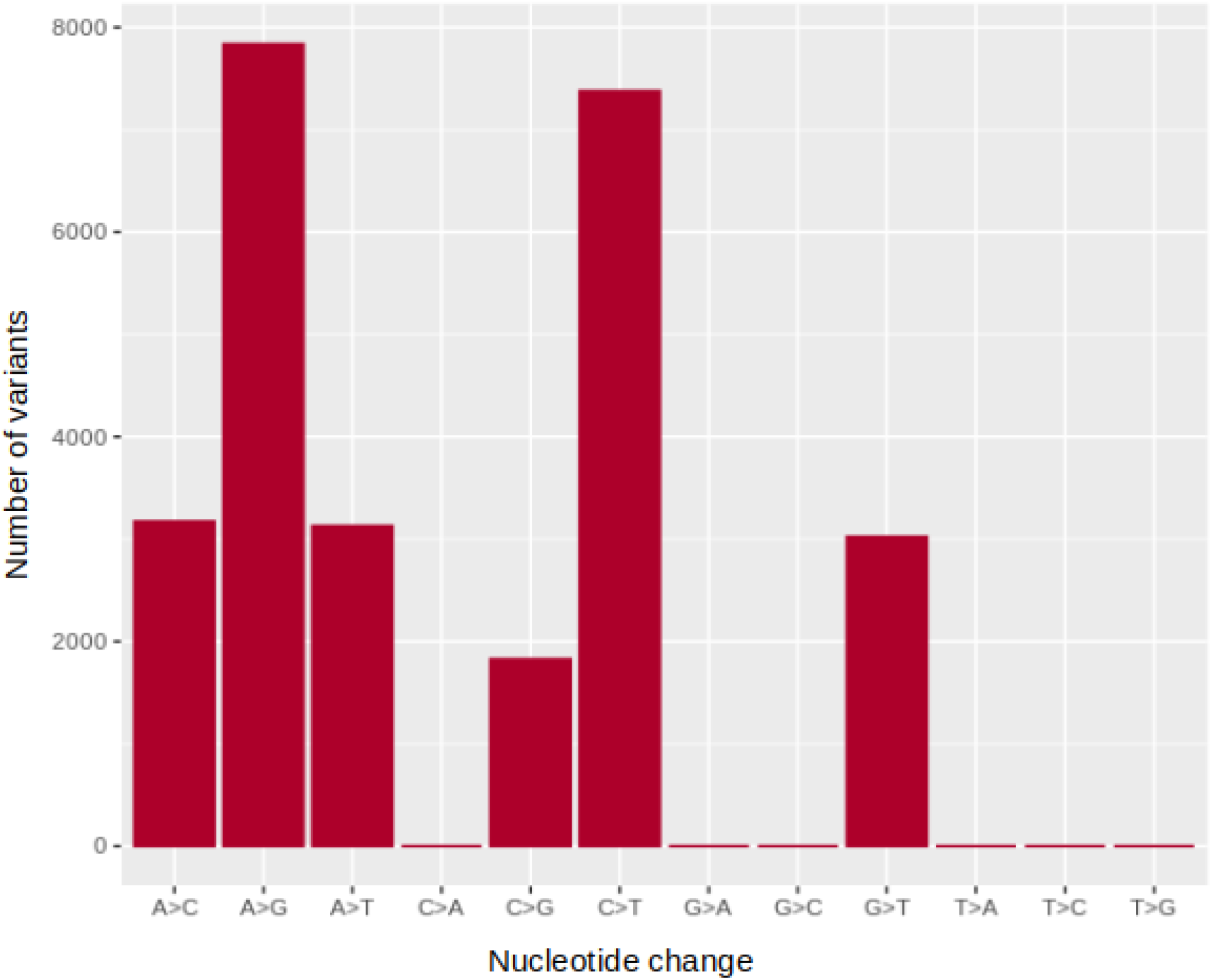
Predicted heterozygous SNP variants by nucleotide change.

## Summary

The methods described here are well suited to any laboratory wishing to reduce expenditures on routine assays and equipment. They are particularly suited for those with limited infrastructure. In addition, the described platform would also be ideal for a teaching module that integrates a fun DIY component to de-mystify molecular methods as well as teach basic genomics and bioinformatics. The inherent low cost of these methods will therefore appeal to budget-minded teaching labs, and can be used to analyze plants in botanical collections, or in ecological studies that incorporate sampling of wild plants from interesting and remote environments.

## Acknowledgments

We thank Marcos Olivos Trujillo for work in preparing the blue light box. This research was funded by core funding from the Agriaquaculture Nutritional Genomic Center. The authors declare that there is no conflict of interest.

## Author contributions

Conceptualization, RHT, CEO, and BJT; Methodology, RHT and BJT; Investigation, RTH, CEO & BJT; Writing – Original Draft, RHT and BJT; Writing – Review & Editing: RHT, CEO & BJT; Supervision, BJT.

## Notes

https://www.ncbi.nlm.nih.gov/sra/?term=PRJNA613471

### Competing Interest Statement

The authors have declared no competing interest.

## References

1. Bioversity International. Mainstreaming agrobiodiversity in sustainable food systems: Scientific foundations for an agrobiodiversity index [Internet]. Bioversity International. Available from: https://cgspace.cgiar.org/handle/10568/89049.

2. Buckler ES, Thornsberry JM, Kresovich S. Molecular diversity, structure and domestication of grasses. Genet. Res. 77(3), 213–218 (2001).

3. Mascher M, Schreiber M, Scholz U, Graner A, Reif JC, Stein N. Genebank genomics bridges the gap between the conservation of crop diversity and plant breeding. Nat. Genet. 51(7), 1076–1081 (2019).

4. Tanksley SD, McCouch SR. Seed banks and molecular maps: unlocking genetic potential from the wild. Science. 277(5329), 1063–1066 (1997).

5. FAO (Food and Agriculture Organization). Women - users, preservers and managers of agrobiodiversity [Internet]. FAO, Rome (1999). Available from: http://citeseerx.ist.psu.edu/viewdoc/download;jsessionid=2BB791DFD15ED4EF10EAE1AC83D930E3?doi=10.1.1.395.2601&rep=rep1&type=pdf.

6. Fernie AR, Yan J. De Novo Domestication: An Alternative Route toward New Crops for the Future. Molecular Plant. 12(5), 615–631 (2019).

7. Till BJ, Reynolds SH, Weil C, et al. Discovery of induced point mutations in maize genes by TILLING. BMC Plant Biol. 4, 12 (2004).

8. Xin Z, Chen J. A high throughput DNA extraction method with high yield and quality. Plant Methods. 8, 26 (2012).

9. Chase MW, Hills HH. Silica Gel: An Ideal Material for Field Preservation of Leaf Samples for DNA Studies. Taxon. 40(2), 215–220 (1991).

10. Liston A, Rieseberg LH, Adams RP, Do N, Ge-lin Z. A Method for Collecting Dried Plant Specimens for DNA and Isozyme Analyses, and the Results of a Field Test in Xinjiang, China. Annals of the Missouri Botanical Garden. 77(4), 859–863 (1990).

11. Till BJ, Jankowicz-Cieslak J, Huynh OA, Beshir MM, Laport RG, Hofinger BJ. Low-Cost Methods for Molecular Characterization of Mutant Plants: Tissue Desiccation, DNA Extraction and Mutation Discovery: Protocols [Internet]. Springer Nature. Available from: http://library.oapen.org/handle/20.500.12657/27721.

12. Hussain M, Iqbal MA, Till BJ, Rahman M-. Identification of induced mutations in hexaploid wheat genome using exome capture assay. PLOS ONE. 13(8), e0201918 (2018).

13. Li G, Jain R, Chern M, et al. The Sequences of 1504 Mutants in the Model Rice Variety Kitaake Facilitate Rapid Functional Genomic Studies. The Plant Cell. 29(6), 1218–1231 (2017).

14. Huynh OA, Jankowicz-Cieslak J, Saraye B, Hofinger B, Till BJ. Low-Cost Methods for DNA Extraction and Quantification [Internet]. In: Biotechnologies for Plant Mutation Breeding: Protocols. Jankowicz-Cieslak J, Tai TH, Kumlehn J, Till BJ (Eds.), Springer International Publishing, Cham, 227–239 (2017) [cited 2019 Oct 2]. Available from: https://doi.org/10.1007/978-3-319-45021-6_14.

